# A Multithreaded Model for Cancer Tissue Heterogeneity: An Application

**DOI:** 10.1101/2022.09.05.505544

**Authors:** Anik Chaudhuri, Shabnam Choudhury, Anwoy Kumar Mohanty, Manoranjan Satpathy, M. Shell, J. Doe

## Abstract

Studying the heterogeneity in cancerous tissue is challenging in cancer research. It is vital to process the real-world data efficiently to understand the heterogeneous nature of cancer tissue. GPU compatible models, which can estimate the subpopulation of cancerous tissue, are fast if the size of input data, i.e., the number of qPCR (quantitative polymerase chain reaction) gene expression reading is extensive. In the real world, we rarely get that much data to reap the benefits of a GPU’s parallelism. Real experimental data from fibroblasts are much less, and models using those data on a GPU are slower than the CPU multithreaded application. This paper will show a method to run GPU-compatible models for cancer tissue heterogeneity on a multithreaded CPU. Further, we also show that the model running on a multithreaded CPU is faster than the model running on a GPU with real experimental data.

## 1. Introduction

TUMOR cells are heterogeneous [1] and [2]. The clonal evolution models suggest that tumor cells accumulate mutation as it progresses. This stepwise accumulation results in various sub-population in a tumor cell and makes it heterogeneous. Another dominant theory called the stem cell suggests that only a small portion of the tumor cell are dominant [3], [4], [5] and [6]. These theories suggest that cancer tissues are heterogeneous and pose specific challenges in cancer treatment. The sub-population reacts differently to a given therapy. It may also happen that a particular combination of drugs works for a patient but may not work for another patient with a different combination of sub-population. It is essential to know the proportion of these sub-populations to make appropriate decisions about therapy. A mathematical model that can account for the heterogeneous behavior of cancer tissue can provide better insight into cancer treatment. Therefore, it is imperative to model the heterogeneity of cancer tissue mathematically.

Authors in [7] discussed a hierarchical model to analyze cancer tissue heterogeneity. The authors in [7], used a Boolean network collection to model cancer, and the weights of each of those networks represented the proportion of each heterogeneous sub-population. A hierarchical model was used to demonstrate the relationship between gene expression measurements and the unknown parameters. In [8], the authors presented a parallelizable model to analyze cancer tissue heterogeneity. Unfortunately, this parallelizable model, which is compatible with a GPU’s SIMD (single instruction multiple data) architecture, does not perform well for a small dataset. The parallelizable model running on a GPU is suitable with large synthetic data but not for real experimental data because the amount of real experimental data is significantly less. So, a GPU is not an ideal choice for such cases. This paper describes a method to run parallelizable models on a multithreaded CPU. We shall show that a multithreaded CPU’s run time is less than the GPU for real experimental data.

## II. MOTIVATION

Relations between proteins and DNA is responsible for cellular interaction [2] and [9]. Using gene regulatory networks is an excellent way to model cell behavior and develop better therapy. Gene regulatory networks have been modeled by differential equation [10], deterministic and probabilistic Boolean network [9], [11] and Bayesian and Dynamic networks [12] and [13]. It is difficult to use probabilistic Bayesian networks to learn parameters from real data due to the huge search space of the parameters.

Authors in [14] and [15] modeled cancer as a “stuck-at” fault in the Boolean network. Faulty logic gates represented the faulty molecules in the transduction network. Much information about cellular interaction is available in the literature; authors in [14] used this prior knowledge to generate a Boolean network from pathway knowledge using the Karnaugh map. The authors produced a Boolean network for the Mitogen-Activated Protein Kinase (MAPK) transduction pathway with this method.

In [7], and [8], authors used a combination of these Boolean networks to estimate the proportion of each subpopulation. The weights of the Boolean network,i.e., the subpopulations, were evaluated by Markov Chain Monte Carlo (MCMC) methods. The model in [7] is not parallelizable, so its computation time increases with an increase in data. The model in [8] is parallelizable, and its computation time does not increase with an increase in data as long as hardware resources are not exhausted. Still, this model on a GPU is slow if the data set size is very small. Real-world experimental data is very small, so running the parallelizable on a GPU is not the best solution. Models running on a CPU are faster than a GPU with a small dataset. In this paper, we shall show a way to run a parallelizable model on a multithreaded CPU to overcome the problem faced by a GPU with a small dataset.

## III. METHOD

The goal is to estimate the proportion of the subpopulations from gene expression data. The Boolean networks are used to model each subpopulation, and the weight exerted on each subpopulation on the observables is estimated. A reasonable way to model gene expression is modelling it with a Normal distribution [7]. In [7] and [8], the probability distribution of the gene expression ration depends on the weight of each Boolean network *α*_*i*_, a coefficient of variation *c* and expression profile *d*_*i*_. *d*_*i*_ is a vector of length *N*, where *N* is the number of subpopulations, i.e., the number of Boolean networks. Here, *i* represents the *i*^*th*^ gene. The expression profile represents the transcription of the observed gene. A value of 1 in the expression profile vector represents an upregulated gene and a value of 0 represents a downregulated gene for a given stimuli.

### A. GPU compatible model on a multithreaded CPU

Figure 1 shows the Bayesian network of our probability model. In this model, there are *V* genes, i.e, *i* ranges from 1 to *V*. Each weight vector *α*_*i*_ is associated with with one gene expression reading *r*_*i*_ for the *i*^*th*^ gene. Here, *α*_*i*_s depend on the vector *K*, and the gene expression measurement *r*_*i*_ depends on *α*_*i*_ and the coefficient of variation *c*. Each *α*_*i*_ is a vector of *N* elements, here, *N* represents the number of subpopulation or the number of Boolean networks. All the elements of *α*_*i*_ should sum up to one.

**Fig. 1:**
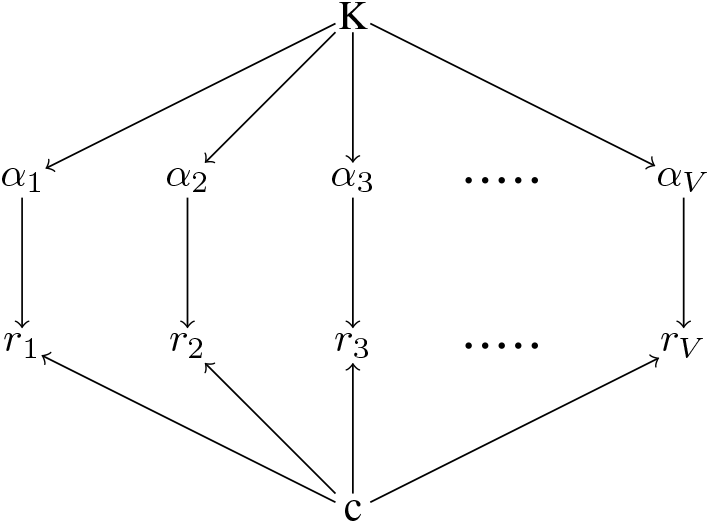
The Bayesian network used in [8].

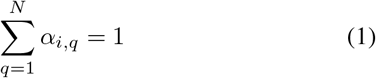

Since, *r*_*i*_ ranges from *r*_1_ to *r*_*V*_, so *α*_*i*_ ranges from *α*_1_ to *α*_*V*_.

The probability distribution of the normalized gene expression ratio *r*_*i*_ for the *i*^*th*^ gene from [8] is

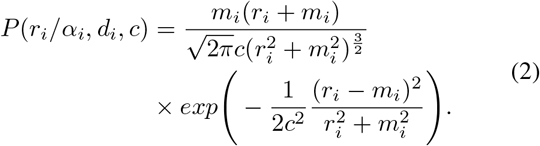

Here, 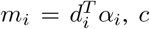, is the coefficient of variation, *r*_*i*_ is the gene expression ratio. The *α*_*i*_ for the *i*^*th*^ gene is

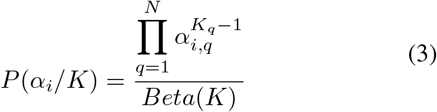

where *Beta (K)* is

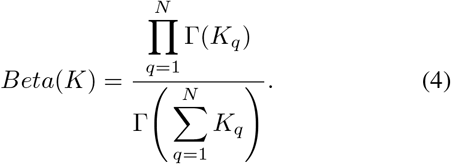

Here, Γ is a Gamma function. *K* and *c* are the unknown parameters which is estimated by using a MCMC (Markov Chain Monte Carlo) algorithm called M-H (Metropolis-Hastings). M-H is an algorithm to sample from an unknown distribution [16], here the posterior distribution of *k* and *α*_*i*_ is unknown, so the M-H algorithm is used.

To calculate the posterior of the unknown, i.e., *K* and *c*, prior distributions are set. The prior over 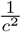 is a Gamma distribution

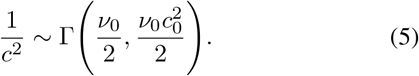

Here, Γ is a Gamma distribution. The prior over *K* is an exponential distribution. The means of this distribution are well separated.

The full conditional of *α*_*i*_ is

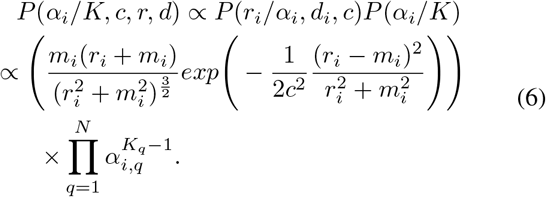

Considering *P* (*K*) as the prior distribution over *K*.

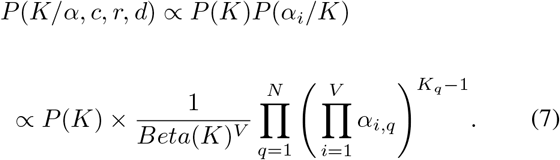

The full conditional of *c* is:

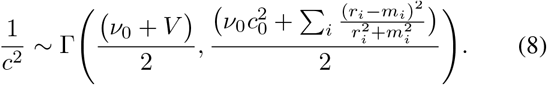

Since the distribution of *K* and *α*_*i*_ are unknown so, Random Walk Metropolis Hastings will be used to sample from these distributions. Figure 2 explains the sampling from the full conditional of *α*_*i*_ and *K* on a multithreaded CPU.

**Fig. 2:**
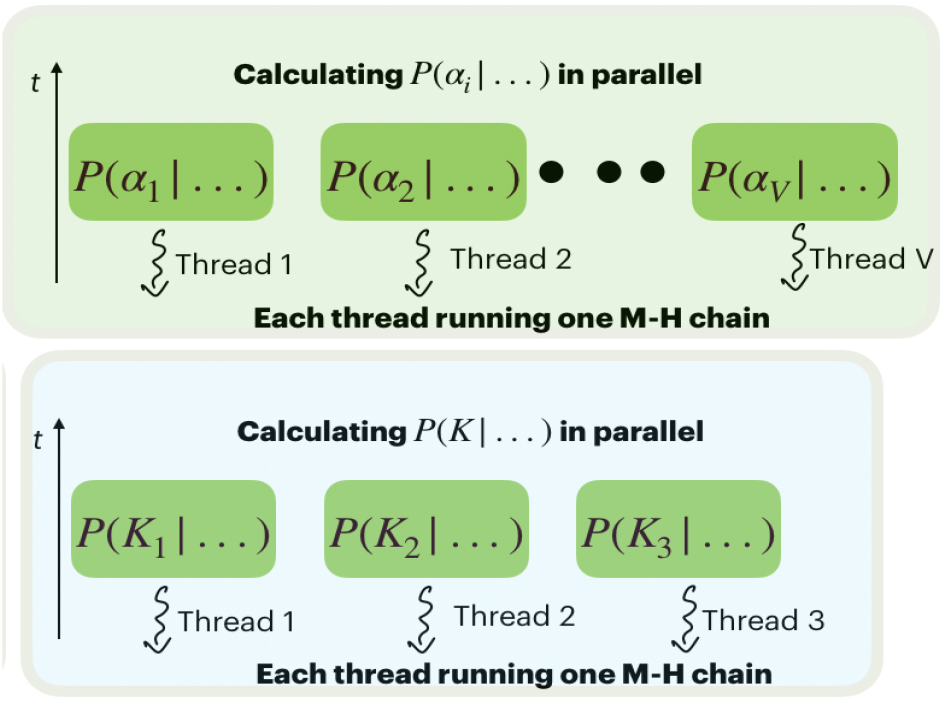
Sampling of equation 6 and 7 on a multithreaded CPU.

Considering each *α*_*i*_ has three subpopulation,i.e., *N* = 3. Letting *V* be the number of genes. The proposal distribution for 6 is a Dirichlet distribution with parameter 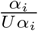. The acceptance ratio is

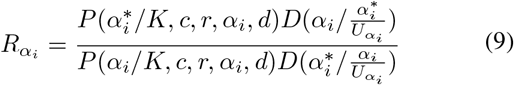

M-H algorithm is used to accept new proposals 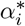.

The proposals 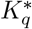 are sampled from a uniform distribution. The acceptance ratio is

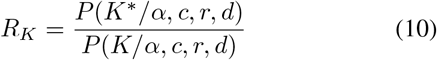

### B. Experiments with synthetic data

The algorithm was written in OpenMP and run until convergence was achieved. The synthetic data was generated by considering

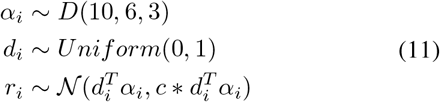

Here, *α*_*i*_ is Dirichlet distributed with parameter *K*, which has been fixed to (10 6 3). *r*_*i*_ is Normally distributed and *d*_*i*_ is uniform distributed.

The unknown parameters are sampled on a multithreaded CPU as shown in figure 2. Metropolis-Hastings algorithm was used to sample from the unknown distributions. The Markov chain was reached stationary at 3000 iterations, but it was run for 10000 iterations to be sure. The estimates of *K* came out to be (10.6 6.1 2.8)^*T*^, these results are very close to the actual value (10 6 3)^*T*^. The *α*_*i*_s are sampled from a Dirichlet distribution by considering the *K*s as the parameters of the distribution, the modes of the *α*_*i*_ are (0.5145 0.3743 0.1112)^*T*^ which is very close to the actual values (0.5165 0.3746 0.1089)^*T*^.

Figure 3 shows the CPU and GPU runtime. The figure shows that for multithreaded CPU code is faster than the single threaded CPU code and the GPU code for data size less than 300.

**Fig. 3:**
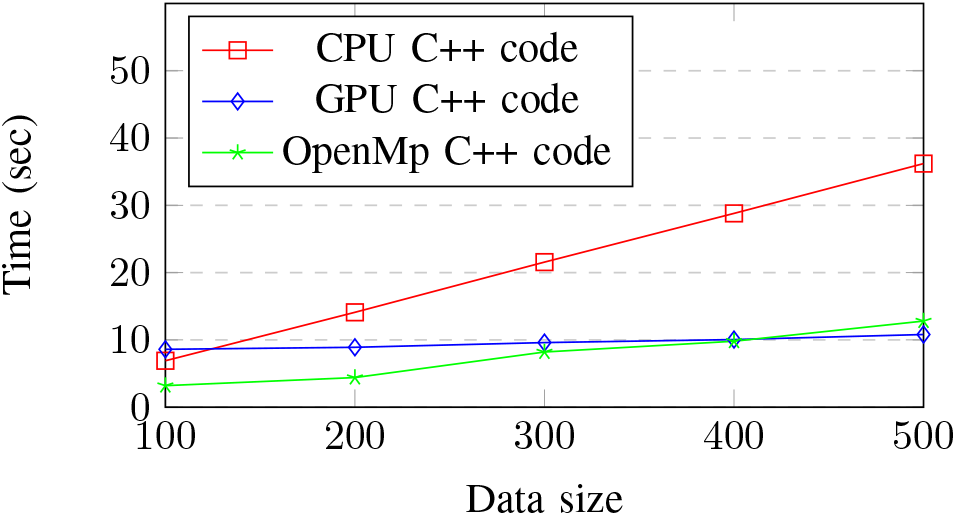
Comparision of CPU and GPU Run Time for 10,000 Monte Carlo iterations

## IV. Experiments with real data

To check the correctness of the model, the algorithm was run with real world experimental data collected from [7]. Samples are drawn from the posterior distribution of *K* using the M-H algorithm; the previous section discussed the sampling procedure. The marginal distribution of the three components of *α* are estimated as described before. The modes of the distribution is (0.6161 0.3236 0.0603)^*T*^, which is very close to the results obtained in [7], i.e., (0.64530.22550.1292)^*T*^. The faultless network has the maximum influence on the observables.

## V. Conclusion

This paper addresses an important problem of designing an algorithm that can be parallelized to study cancer tissue heterogeneity. This algorithm uses prior pathway knowledge to estimate the proportion of each subpopulation. The gene expression was modeled as ratios of normally distributed random variables whose means are affected by the networks included. We also demonstrated how to use M-H MCMC algorithm on a multithreaded CPU to estimate the unknown parameters. This estimate is useful to find out the dominant subpopulation among all the subpopulations. We used the algorithm in [18] to sample from a Gamma distribution on a GPU. This helped us to parallelize the algorithm and reduce the computation time.

